# An Efficient Parallel Sketch-based Algorithmic Workflow for Mapping Long Reads

**DOI:** 10.1101/2023.11.28.569084

**Authors:** Tazin Rahman, Oieswarya Bhowmik, Ananth Kalyanaraman

## Abstract

Long read technologies are continuing to evolve at a rapid pace, with the latest of the high fidelity technologies delivering reads over 10Kbp with high accuracy (99.9%). Classical long read assemblers produce assemblies directly from long reads. Hybrid assembly workflows provide a way to combine partially constructed assemblies (or contigs) with newly sequenced long reads in order to generate improved and near-complete genomic scaffolds. Under either setting, the main computational bottleneck is the step of mapping the long reads—against other long reads or pre-constructed contigs. While many tools implement the mapping step through alignments and overlap computations, alignment-free approaches have the benefit of scaling in performance. Designing a scalable alignment-free mapping tool while maintaining the accuracy of mapping (precision and recall) is a significant challenge. In this paper, we visit the generic problem of mapping long reads to a database of subject sequences, in a fast and accurate manner. More specifically, we present an efficient parallel algorithmic workflow, called `JEM-mapper`, that uses a new minimizer-based Jaccard estimator (or JEM) sketch to perform alignment-free mapping of long reads. For implementation and evaluation, we consider two application settings: (i) the hybrid scaffolding setting, where the goal is to map a large collection of long reads to a large collection of partially constructed assemblies or contigs; and (ii) the classical long read assembly setting, where the goal is to map long reads to one another to identify overlapping long reads. Our algorithms and implementations are designed for execution on distributed memory parallel machines. Experimental evaluation shows that our parallel algorithm is highly effective in producing high-quality mapping while significantly improving the time to solution compared to state-of-the-art mapping tools. For instance, in the hybrid setting for a large genome *Betta splendens* (≈350*Mbp* genome) with 429*K* HiFi long reads and 98*K* contigs, `JEM-mapper` produces a mapping with 99.41% precision and 97.91% recall, while yielding 6.9`×` speedup over a state-of-the-art mapper.

## 2 Introduction

Over the last two decades, numerous genomes have been assembled using short read sequencing technologies. These technologies continue to present a cost-effective and high-throughput solution to sequencing. While short reads are accurate (*<* 1% error) the challenge is in their lengths (100 to 250 bp), which causes fragmented assemblies of contigs (≈ 10^3^ − 10^4^ bp) that are several orders of magnitude shorter than their target genomes (≈ 10^6^ − 10^9^). Recent emergence in long read sequencing technologies represents a significant advance in addressing this challenge. The first generation of long read technologies (e.g., PacBio SMRT [1] or Oxford Nanopore ONT [2]) produce reads that are over 10 Kbp but also have a larger error rate (between 11%−14% [3]). The more recent generation of technologies such as PacBio HiFi (High Fidelity) [4] have highly improved accuracy (99.9%). There are also several long read error correction tools [5]. Given these, the prospect of assembling long contiguous portions of the genome has dramatically improved [6].

Broadly speaking, two classes of approaches exist for using long reads—standalone and hybrid. *Standalone* long read assemblers [6, 7] produce a *de novo* assembly from long reads using the Overlap-Layout-Consensus (OLC) paradigm [8]. HiFi long reads significantly facilitate the task of *standalone* assemblers and with higher accuracy in the reads, the overlap graph becomes smaller as compared to the overlap graphs generated from error-prone long reads. However, the OLC paradigm requires pairwise alignments of long reads which is the major computational bottleneck of *standalone* assemblers.

The computational burden is exacerbated by the fact that more sequencing coverage is needed for *de novo* sequencing. For instance, this step took about 95% of the time while assembling *D. melanogaster* using [9].

*Hybrid assemblers* [10, 11], on the other hand, offer the benefit of combining long and short reads, or alternatively, combining long reads with prior constructed assemblies from short reads (i.e., contigs). The use of prior constructed contigs (in lieu of short reads) can improve scalability since the number of contigs tends to be orders of magnitude fewer than the number of short reads. By combining long reads with contigs we aim to extend the contigs into longer scaffolds through contig-to-contig linking information that may be available in the long reads. Depending on the genomic fraction covered by the contigs, this may also imply that it is possible to produce long scaffolds with a decreased sequencing coverage (and cost) in long read sequencing (compared to a *de novo* pipeline). In order to implement this hybrid strategy, however, we need a way to efficiently map the long reads to the contigs.

These two cases of mapping long reads motivate the development of the mapping strategy proposed in this paper. Fig. 1 illustrates these two mapping cases. Fig. 1(a) shows the case of mapping long reads for the hybrid setting, where the goal is to map long reads (shown as queries 𝒬) to partially constructed assemblies or contigs (considered subjects 𝒮). The information computed by mapping can be used by a downstream scaffolder to fill in the assembly gaps between adjacent contigs and generate longer scaffolds. Fig. 1(b) shows the case of mapping long reads for the *de novo* long read assembly setting, where the goal is to map the long reads against themselves so as to detect overlapping long reads. Note that in both cases the reference genome 𝒢 is assumed to be unknown. As a convention henceforth, we refer to the mapping application for the hybrid setting as `L2C` and the mapping application for the *de novo* assembly setting as `L2L`.

**Figure 1.**
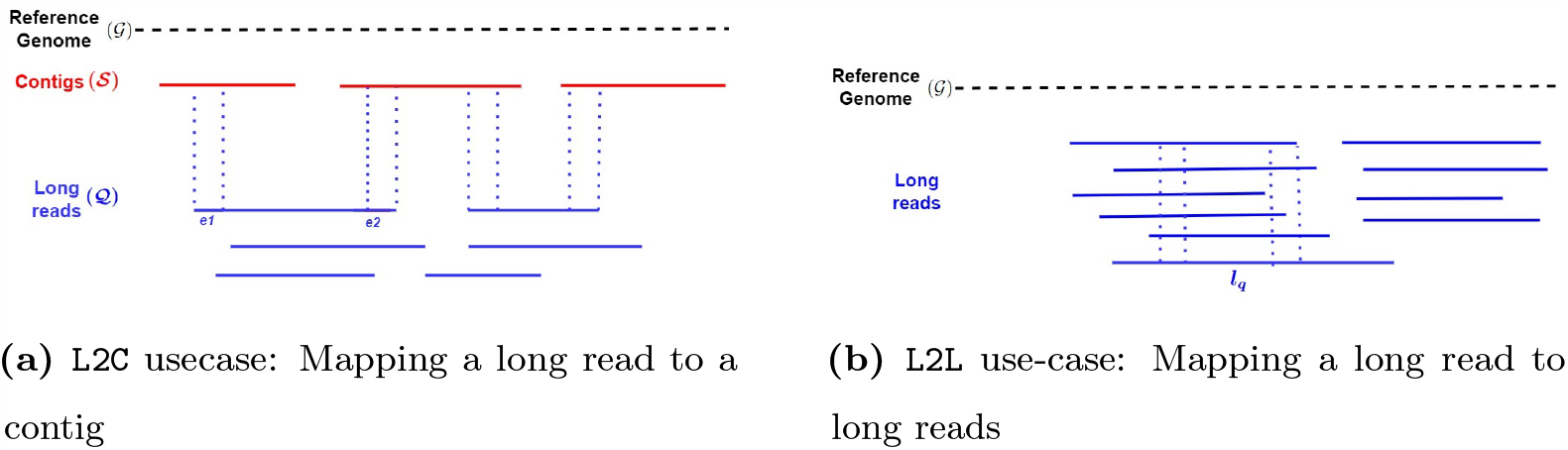
Our target use-cases: a) `L2C` mapping use-case: Subjects are a set of contigs and queries are a set of long reads. Each end of a long read can be expected to map to a single contig (assuming a non-redundant contig set). b) `L2L` mapping use-case: Subjects and queries are a set of long reads. For each long read *l*_*q*_, the goal is to find its set of overlapping long reads. The reference genome (𝒢) is assumed to be unknown, and is only shown to display the mapping coordinates.

While there are a number of sequence mapping approaches (as reviewed in §3), scalability of these tools and in addition their ability to work with different types of sequences (contigs, long reads) are limitations. The key to performance scalability is to reduce alignment computations between long reads and the corresponding subjects.

However, alignment-free approaches could suffer low precision and/or recall (see §3).

**Contributions** In this paper, we present a new parallel algorithmic workflow for fast and accurate mapping of long reads, under both `L2C` and `L2L` use-cases. More generically, the inputs are a set of queries 𝒬 and a set of subjects 𝒮. The output is a mapping for each *q* ∈𝒬 (as formally defined in §4). The main contributions of the paper are as follows.

- Methods: We present a new sketching-based method for alignment-free mapping of long reads. As part of our approach, we propose a minimizer-based Jaccard estimator (or JEM) sketch, that is a variant of the classical MinHashing (§4.2).
- Algorithmic workflow: We present an efficient and scalable parallel algorithmic workflow that uses the JEM sketch to perform mapping of long reads on distributed memory parallel machines. We adapt this workflow to provide two parallel implementations, one for `L2C` and another for `L2L`.
- Evaluation: We conduct a thorough empirical evaluation of the proposed sketching-based implementations for both of the use-cases. Results show that our method is able to match the mapping quality of a state-of-the-art mapping tool, while providing significant speedups over the state-of-the-art. In particular, for the `L2C` use-case, our distributed memory implementation running on 64 processes achieves speedups between 6.9× to 13× compared to the state-of-the-art multithreaded execution on 64 threads. Similarly, for the `L2L` use-case, for the complex genomes, our implementation running on 64 processes achieves speedups between 7.6× to 15× compared to the state-of-the-art multithreaded execution on 64 threads.

We refer to the newly proposed algorithmic workflow as `JEM-mapper`, and the source code is available as open source in https://github.com/TazinRahman1105050/JEM-Mapper for testing and application. A preliminary version of our paper appeared in [12].

With mapping applications increasingly becoming more heterogeneous in their data sources, including in hybrid scaffolding/assembly workflows or reference-guided assembly workflows [13], the techniques described in this paper have broad applicability. In what follows, we provide a review of the relevant sequence mapping literature (§3), our parallel approach and algorithmic workflow (§4), and the experimental results and evaluation (§5).

## 3 Related work

Sequence mapping is a classical problem in bioinformatics. It can be abstracted as one of mapping a set of *query* sequences (e.g., reads) to a set of *subject* sequences. Traditional sequencing mapping tools (e.g., [14, 15]) focus on aligning short reads (queries) against a reference genome (subject). The hybrid setting differs from this classical setting in a couple of different ways. First, *in lieu* of the reference (which is typically a handful of very long subject sequences), the input subjects consist of a set of contigs which represent a highly fragmented view of the reference genome.

Consequently, the contig sets can have tens to hundreds of thousands of sequences, and may also widely vary in their sequence lengths (10^3^-10^5^ bp).

As for the query set, long reads are significantly longer than the short reads used in conventional reference mapping. In the absence of more scalable tools, the current batch of hybrid assemblers [10, 11, 16] rely on a classical mapping tool to implement their mapping step. For instance, Haslr [11] first assembles the short reads using Minia [17], and then maps the contigs to the set of long reads using Minimap2 [15]. Similarly, SAMBA [16] maps the long reads to the set of contigs using Minimap2 [15]. We show in the results section (§5) that using a generic mapping tool such as Minimap2 for `L2C` or `L2L` applications results could result in a loss in precision for larger inputs.

The step to compute overlapping pairs of long reads within long read assemblers can also be considered another form of mapping. State-of-the-art long read assemblers depend on either alignment tools (e.g., DALIGNER [20], BLASR [10]) or overlap candidate detection (alignment-free) tools (for example, Minimap2 [15], MECAT [18], or MHAP [6]).

To improve scalability of mapping, there has been a growing interest in alignment-free approaches [21–25], and in particular sketching—e.g., minimizers [25, 26] and MinHashing [27]. Sketching is a class of techniques that use samples derived from the input sets (or sequences) to be compared in order to approximate similarity.

Introduced originally for document clustering [27], sketching and its relatives like minimizers [26] have found extensive use in bioinformatics. While these techniques have shown significant promise for mapping in the classical setting, they have not yet been fully demonstrated for the use-cases targeted in this paper (`L2C` and `L2L`). Among the alignment-free approaches for overlap detection in standalone assemblers, MHAP [6] uses Jaccard similarity between long reads to estimate overlap between sequences.

However, this approach achieves a very low F1 score [8]. So, the balance between precision and recall is not maintained for complex genomes.

Table 1 provides a high-level summary of the different mapping tools that can handle long reads. The mapping tool presented in this paper (`JEM-mapper`) is shown as the last row. It can be used for both `L2C` and `L2L` use-cases, and is the only tool with support for distributed memory parallelism.

**Table 1.** State-of-the-art comparison of relevant sequence mapping tools. The first three columns indicate the type of mapping targeted with the long reads (i.e., the problem scope). The remaining columns on the right refer to the type of approach used.

| Mapping tool | Problem scope |  |  | Approach |  |  |
| --- | --- | --- | --- | --- | --- | --- |
|  | Supports reads to reference genome mapping | Supports reads to contig mapping (L2C) | Supports long read mapping (L2L) | Alignment-based | Sketching-based | Type of parallelism |
| BLASR [10] | ✓ | ✓ | ✓ | ✓ | × | shared-memory |
| MHAP [6] | × | × | ✓ | × | ✓ | shared-memory |
| MECAT [18] | ✓ | × | ✓ | × | × | shared-memory |
| Mashmap [19] | ✓ | ✓ | × | × | ✓ | shared-memory |
| Minimap2 [15] | ✓ | ✓ | ✓ | × | × | shared-memory |
| JEM-mapper (this paper) | × | ✓ | ✓ | × | ✓ | distributed-memory |

## 4 Methods

In this paper, we consider two closely related mapping problem abstractions, motivated by two specific use-cases, as described below. Figure 1 illustrates the two use-cases. Let 𝒬 denote a set of query sequences, and 𝒮 denote a set of subject sequences. The sequences use an alphabet Σ (DNA: Σ = {a, c, g, t}); therefore, 𝒬⊆ Σ^*^ and 𝒮⊆ Σ^*^. Let *ψ* : Σ^*^ × Σ^*^ → ℝ_≥0_ be a mapping function to map a query *q* to a subject *s*.

### Definition 1.

**The** `L2C` **mapping use-case:** *Given* 𝒬 *and* 𝒮, *find for each query q* ∈ 𝒬 *a best mapping subject* 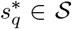, *given by*,

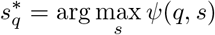

Setting 𝒬 to a set of input long reads and 𝒮 to a set of input contigs (hence the term, `L2C`) would make the results of mapping useful in a hybrid scaffolding workflow.

### Definition 2.

**The** `L2L` **mapping use-case:** *Given* 𝒬 *and* 𝒮, *and given a mapping quality threshold τ*, find *for each query q* ∈ 𝒬 *the set of subject sequences A*_*q*_ ⊆ 𝒮 *such that:*

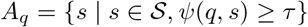

Setting 𝒬 and 𝒮, both to a set of input long reads (hence the term, `L2L`), would help identify overlapping pairs of reads for a long read assembler.

While the function *ψ* can be implemented as a sequence alignment, computing alignments at scale can be expensive. Therefore, alignment-free approaches are more desirable in practice. Our approach uses an alignment-free sketch to reduce the search space as described below.

### 4.1 Preliminaries and notation

`MinHash` **preliminaries** The `MinHash` sketching scheme was introduced by Broder in 1997 [27], originally to compute resemblance or Jaccard similarity between documents. Given two sets *A* and *B*, the Jaccard similarity between the sets is given by: 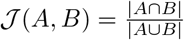. In this seminal work, Broder showed that there exists a family of permutations (*π* : [*n*] → [*n*]) called the minwise independent permutations, which can be used to generate fixed size sketches from the sets *A* and *B*. It then suffices to compare the sketches instead of explicitly computing the 𝒥 (*A, B*), i.e.,

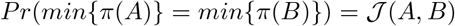

In other words, higher the Jaccard similarity, higher the probability that the sketches obtained *A* and *B* will match. To improve the chance that a random sketch is found, a fixed number of random trials (*T*) is executed. This is achieved by choosing *T* random minwise independently permutations: {*π*_1_, *π*_2_, …, *π*_*T*_ }, and using them to generate the `MinHash` *sketches* for sets *A* and *B*, denoted by 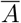 and 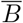 respectively:

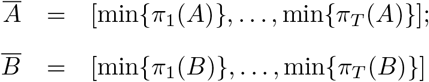

Subsequently, if any of the trials produce the same minimum between 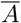 and 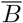 then we conclude *A* is *similar* to *B*. In practice, a value around 100 to 200 is used for *T* [27]. We refer to the `MinHash` sketches (e.g., 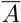, 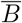) sometimes as just “MinHashes” for the underlying sets.

**String notation** Let *s* denote an arbitrary string over alphabet Σ, and let |*s*| denote its length. We use the terms strings and sequences interchangeably. A *k*-mer is a (sub)string of length *k*. Given Σ and *k*, let 𝒦 denote the set of all *k*-mers that can be built using Σ. Note that |𝒦|=|Σ|^*k*^. We use the term *canonical ordering* of *k*-mers, denoted by 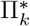, to refer to the lexicographical ordering of the *k*-mers in 𝒦. For instance, if *k*=2, the canonical ordering of 𝒦 is given by: 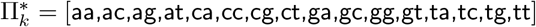. Given a string *s* ∈ Σ^*^ and a choice of *k*, the notation *s*_*k*_ is used to denote the set of all *k*-mers present in *s*.

### 4.2 Computing mapping using a minimizer-based Jaccard estimator sketch

In the mapping applications, we have two sets of strings—𝒬 containing queries, and 𝒮 containing subjects. For a string *s*, its `MinHash` sketch can be constructed from the set of all *k*-mers (*s*_*k*_) in *s*—i.e., during trial *t*, apply a hash function *h*_*t*_(.) on each *k*-mer in *s*_*k*_ and then select the *k*-mer with the minimum value as part of the sketch.

Using this idea, a simple way to apply the MinHashing scheme for `mapping` is as follows. First enumerate the MinHashes (or the sketches) for each subject (one minimum for each random trial *t* ∈ [1, *T* ]) and insert those into a sketch data structure 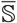. Subsequently, during querying time, sketches are also generated from each query. The more sketches a query generates in common with a subject, the higher the likelihood that it shares a high sequence level similarity. Therefore, we can simply track the frequency of the subjects that “hit” with a given query, and report the set of top matching subjects (if any) as the mapped output hit to that query. This simple algorithmic workflow is illustrated in Fig. 2.

**Figure 2.**
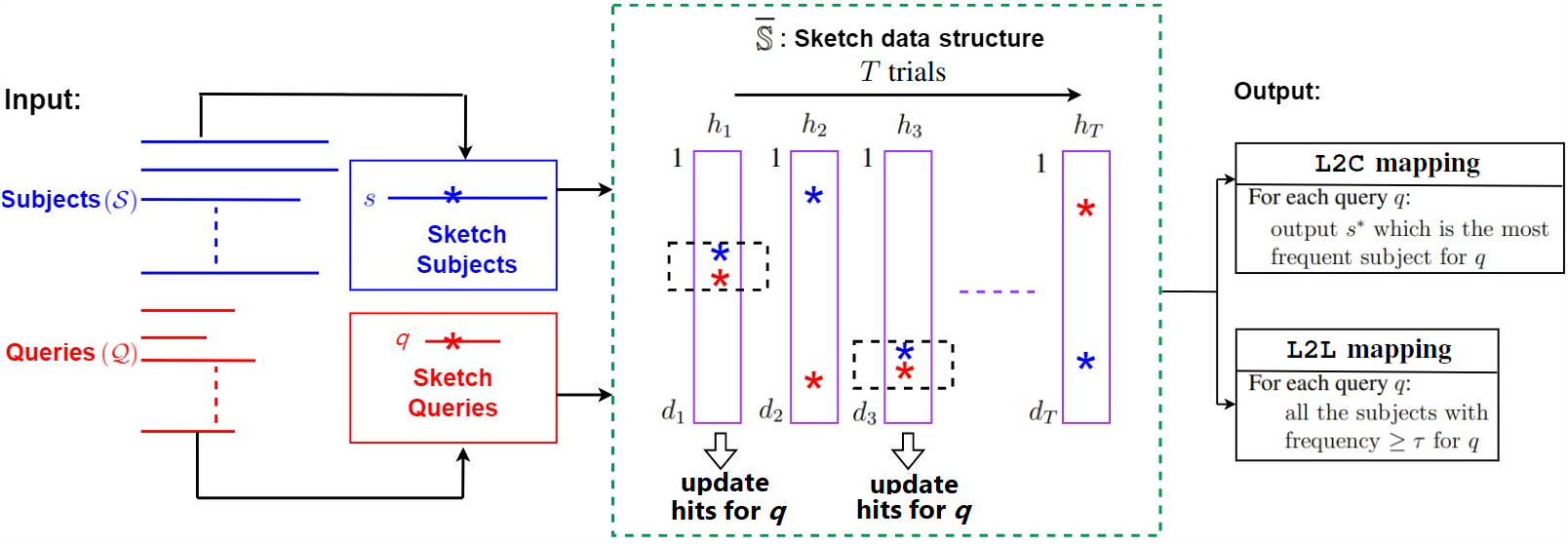
`JEM-mapper`: Illustration of the major steps of our sketch-based mapping algorithm, `JEM-mapper`. The workflow shown is generic to work for both `L2C` and `L2L` use-cases.

While this workflow can be efficiently implemented, we make several changes, as there are a few key challenges with a direct application of MinHashing as described above.

First, note that in the `mapping` applications targeted, subjects and queries could have significantly differing lengths. If a query *q* is significantly longer (say 10*Kbp*) than a corresponding mapped subject *s* (say 3*Kbp*), then even if *s* has significant identity within *q*, MinHashing may select *k*-mers that may lie outside the region of the overlap. This could mean missing out on a true mapped (affects recall). Similarly, if a subject *s* is significantly longer (say, 20*Kbp*) compared to a query *q* (say 10*Kbp*), recall could again be affected as the sketches from *s* could lie outside the region of true overlap. Either way, the qualitative efficacy of `MinHash` for mapping could be negatively impacted.

To overcome this limitation, we use two ideas: a) to map only segments of queries; and b) to compute a minimizer-based Jaccard estimator (instead of the classical `MinHash` form).

#### 4.2.1 Using the *segments* of a query *q*

Instead of extracting sketches from the entire length of a query *q*, our approach uses only several regions (aka. “segments”) of *q*. Specifically, we define a fixed *segment length ℓ*. We then map only *λ segments* of query *q* to subjects and report respective subjects with hits. The main idea is to focus on regions of high overlaps between the query and subject. This not only improves quality, but also reduces work, making the algorithm faster. In both of our applications (`L2C` and `L2L`), we found a value of *λ* = 2 to be sufficient in our experiments. Henceforth, we revise the set of queries 𝒬 to include the two segments of each query—i.e., if there are *m* queries, then 𝒬 consists of 2*m* sequences of length *ℓ* each. The heuristic to pick *segments* for `L2C` or `L2L` is described in (§4.3).Fig. 1 shows the end segments of queries mapped to subjects.

#### 4.2.2 Sketching using minimizer Jaccard estimate

Segments help constrain the regions where sketches are extracted from the queries. However, subjects can be very long and it is possible that the region of overlap between a query and a subject can span anywhere in its length. Therefore, we follow a two-pronged idea by: a) reducing the base set of *k*-mers for Jaccard similarity computation to a set of *minimizers* [26] obtained from subjects, and b) then using a sliding interval of length *ℓ* bp over the list of those minimizers to select one `MinHash` per interval. The list of minhashes so extracted becomes our version of the minimizer-based Jaccard estimator sketch (abbreviated as “JEM” henceforth) of the subject for that trial *t* ∈ [1, *T* ]. Fig. 3 illustrates this procedure using a conceptual example. The detailed algorithm is as follows.

**Figure 3.**
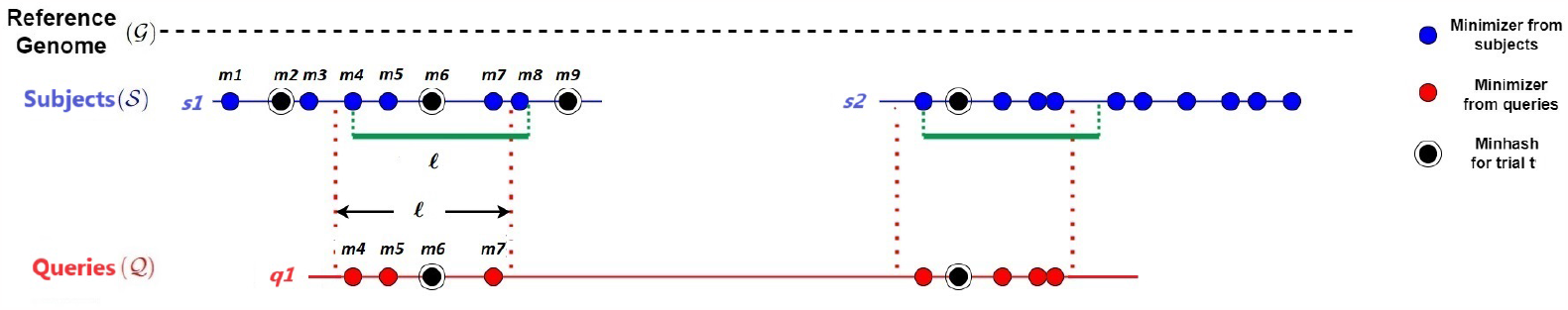
An example to illustrate the way minimizer-based Jaccard estimate sketch (or the JEM sketch) is generated by our approach, `JEM-mapper`. At the subject processing time, the list of minimizer tuples *M*_*o*_(*s, w*) is generated for each contig (shown as blue circles). We then slide an interval of length *ℓ* over the set of minimizers based on their positions. On *s*1, this is shown as the list [*m*_4_ … *m*_8_]. For each such interval, we generate *T* minhashes for *T* trials. The black concentric circle shows the minhash that was randomly selected for trial *t* in that interval (i.e., the sketch). At query processing time, for long read segment, we generate a similar set of minimizers (denoted by the red circles). We then pick *T* JEM sketches in a similar fashion and look for hits in the sketch table before detecting the top hit.

The minimizer-based Jaccard estimate calculates the Jaccard similarity between two sequences using the minimizer sketches between them. Given a sequence *s*, an integer *k* and window size *w*, a minimizer is the *k*-mer with smallest hash value of the *w* consecutive *k*-mers. We use the lexicographically smallest *k*-mer as our hash function, consistent with previous works [28, 29]. The *minimizer sketch* of *s* (denoted by *M* (*s, w*)) is the set of all such minimizers in *s*. Hence the minimizer Jaccard estimate between sequences *A* and *B* is:

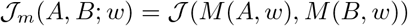

In other words, the minimizer Jaccard estimate allows us to collect and compare sketches from the list of minimizers of a sequence (rather than all the *k*-mers). This reduces work and also provides a certain degree of qualitative robustness against noisy *k*-mers.

Another type of a minimizer based Jaccard estimate has been used prior in the mapping tool Mashmap [21, 30]. Our algorithm is different in the way these sketches are computed. In Mashmap, for each minimizer, a list of all positions it is present in the subject is maintained. Later, during mapping time, if a query shares a minimizer with multiple positions, then the region where the query has maximal local intersection on the subject is detected and reported at query time. This approach entails at first, short-listing positional candidates and then eliminating those that do not have sufficient concentration of query minimizers in their vicinity.

By contrast, our approach directly applies the segment length *ℓ* of the query as the interval length over the subject, and tracks the `MinHash` for each such interval of the subject—as shown in Fig. 3. This guarantees that the sketches are generated at the resolution of the segment length, both for the subjects and queries, thereby obviating the need to check for any distance constraints later.

##### Algorithm 1

*Sketch*_*byJEM*

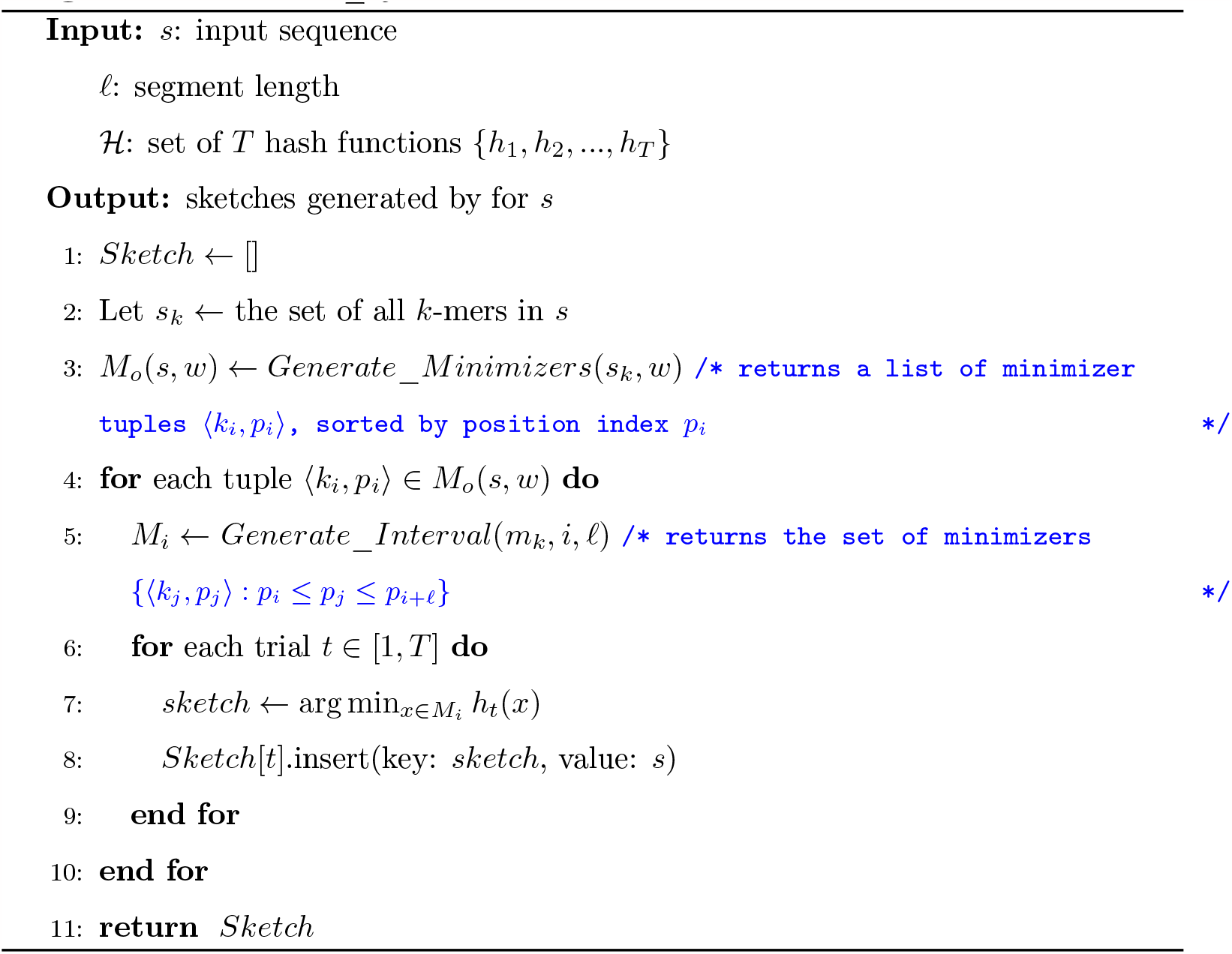

More specifically, let *M*_*o*_(*s, w*) represent the set of all minimizers and their corresponding positions from a string *s*. This is maintained as a set of tuples of the form ⟨*k*_*i*_, *p*_*i*_⟩, where *k*_*i*_ denotes the minimizer at position *p*_*i*_ on *s*. The set *M*_*o*_(*s, w*) is kept sorted based on the minimizer positions. For a given interval length *ℓ*, let us define *M*_*i*_ to be the set of consecutive minimizers in *M*_*o*_(*s, w*) such that *M*_*i*_ = {⟨*k*_*j*_, *p*_*j*_⟩ : *p*_*i*_ ≤ *p*_*j*_ ≤ *p*_*i*_ + *ℓ*}. We slide intervals of length *ℓ* over the set of minimizers and within each interval, *T* minhashes are generated.

Algorithm 1 shows how sketches are extracted using our approach. Algorithm 2 shows the overall `JEM-mapper` algorithm. The algorithm first generates sketches (using Algorithm 1) for all the subjects, and populates them into a sketch data structure 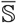. Subsequently, each query is processed by first generating its sketches and looking them up in 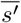. The hits are accumulated with every query minhash colliding with a subject minhash, across the *T* trials. For `L2C`, the subject that has the largest number of hits with a query is reported as the mapped output. For `L2L`, the union of all subject sequences that generated a hit with the query is reported. In our implementation, we use a threshold *τ* for filtering low hit subjects (see the implementation remarks in Section 4.4).

In our implementation, we have used lexicographic ordering of *k*-mers to extract minimizers from window *w*. For a substring *s*^*′*^ of length greater than *k*, a canonical minimizer is the smallest *k*-mer of *s*^*′*^ and its reverse complement 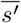 based on lexicographic ordering. To generate the *T* minhashes for each interval, we assign each minimizer of that interval its *k*-mer rank *x* (i.e., as per its canonical ordering in 𝒦), and then use *T* random hash functions of the linear congruential form: *h*_*t*_(*x*) = ((*A*_*t*_ · *x* + *B*_*t*_) mod *P*_*t*_). Subsequently, the *k*-mer corresponding to the smallest hashed value becomes the sketch for that string for trial *t* ∈ [1, *T* ]. Here the triplet ⟨*A*_*t*_, *B*_*t*_, *P*_*t*_⟩ are randomly generated constants associated with the trial *t*. We generate *T* triplets, one for each trial, *a priori*, and use the same *T* triplets to sketch all sequences.

##### Algorithm 2

Mapping by `JEM-mapper` (𝒬, 𝒮)

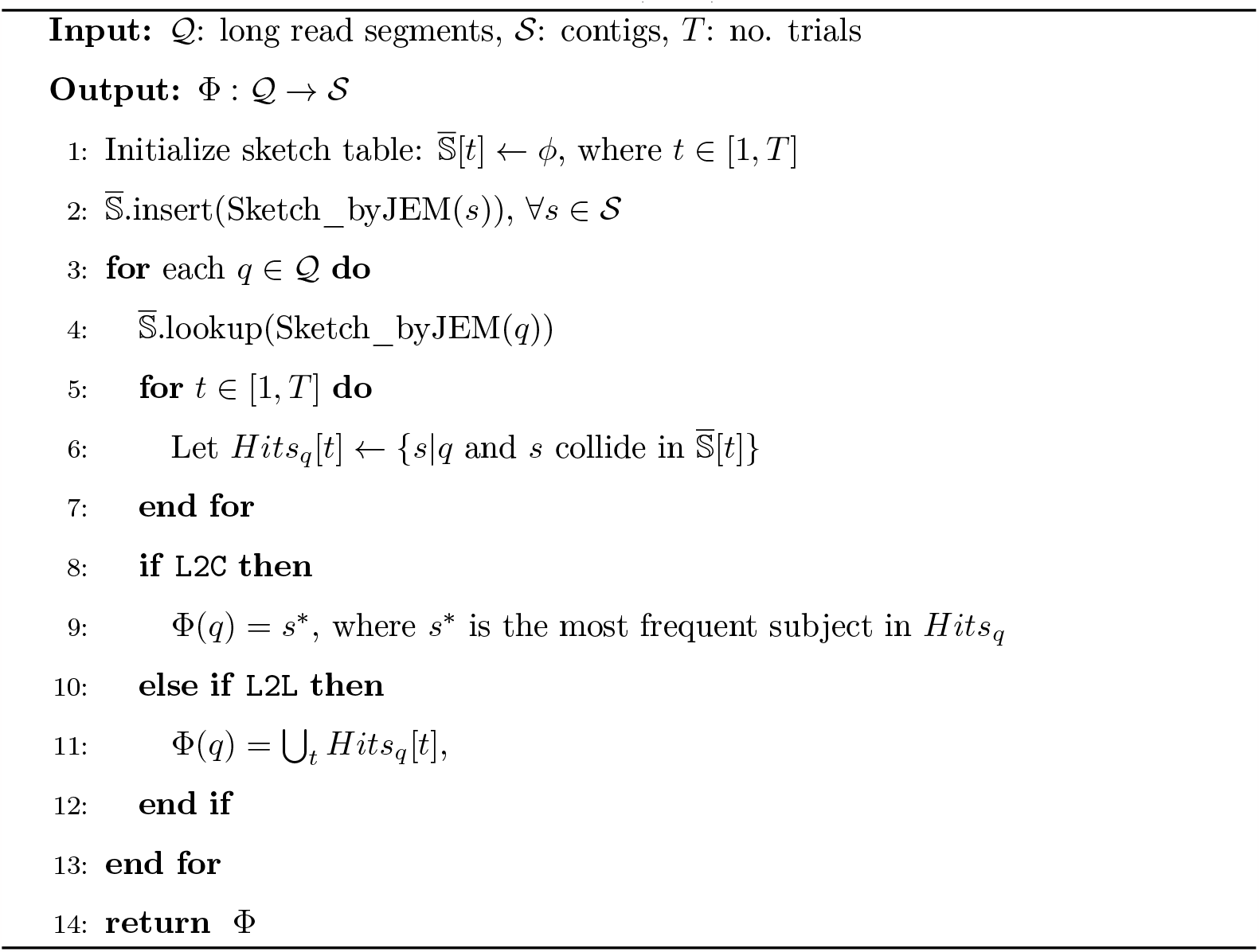

### 4.3 Additional implementation details

#### Masking of repeats in complex input genomes

For complex genomes, particularly for eukaryotic organisms, a significant portion of the whole genome is repetitive. For instance, more than 50% of the human genome is repetitive, with higher fractions for several plant genomes [31–35]. Although long-read sequencing technologies have improved the ability to resolve repeats in genomes, presence of short repetitive stretches can still confound the mapping process. For numerous genomes, repeat information is already available. To take advantage of such available information, we mask the repetitive regions of the input sequences using the RepeatMasker tool [36]. Masked regions appear as ‘N’s. In both `L2C` and `L2L` use-cases, we have used masked input genomes.

#### Segments selection

Since our mapping uses sketches derived from segments, the selection of regions to extract segments could impact the mapping quality. In the case of `L2C`, long reads are mapped to contigs (prior constructed assemblies). Therefore, our implementation extracts segments from the ending regions of a long read—that way, the approach is suited to provide linking information between contigs, i.e., the farthest separated pair of contigs linked by a query long read (as illustrated in Fig. 3). This information can be used by a hybrid scaffolder to increase the span of its scaffold. In our `L2C` experiments, we used a value of *ℓ* = 1000 bp.

For the `L2L` mapping use-case, note that each long read query can map to an arbitrary number of other long read subjects, and the regions of overlap are not constrained to ends of a long read. Therefore, our segment selection scheme does *not* constrain the selection to ends of a long read. Instead we select the top two segments (each of length up to *ℓ* bp) that are non-overlapping on the long read with least ‘N’ content. The rationale for choosing a region with least ‘N’ content is simple—to improve the mapping quality (recall, in particular) for long reads that have repetitive content. We use *ℓ* = 2000 bp as our segment length for `L2L`, based on prior works that have suggested similar lengths [8, 37].

#### Frequency-based heuristics for unmasked inputs

As we have mentioned earlier, we have used lexicographic ordering of *k*-mers to extract minimizers from window *w*. For genomes where the libraries of repeats are not available, hence we used unmasked inputs, we used frequency based heuristic to extract minimizers. This is based on the simple observation that *k*-mers extracted from repeat regions are more frequent in comparison to the *k*-mers from non-repetitive regions. *k*-mers from repetitive regions increase the computational cost, as these *k*-mers generate many incorrect overlaps. So, we have used the least frequent *k*-mer as the minimizer from window *w*. For a substring *s*^*′*^ of length *w*, the least frequent *k*-mer of *s*^*′*^ and its reverse complement 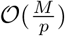 is considered as the minimizer.

### 4.4 Parallelization

We now present a distributed memory parallel algorithm for our `JEM-mapper` algorithm. The Algorithm 2 is well suited for parallelization on a distributed memory as described below. Our implementation is in C/C++ and MPI (for communication). The beta version of this software is available on https://github.com/TazinRahman1105050/JEM-Mapper.

We use the following notation: *m* = |𝒬|; *M* = Σ_*q∈𝒬*_|*q*|; *n* = |𝒮|; *N* = Σ_*s∈𝒮*_ |*s*|; and *p* to denote the number of processes. The major parallel steps are as follows.

S1) *(load input)* The processes load the input 𝒬 and 𝒮 in a distributed manner, such that each process gets approximately 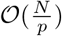 query bases and 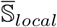 subject bases. Let 𝒬_*local*_ and 𝒮_*local*_ denote the sets of local queries and subjects respectively, held by any given process.

S2) *(sketch subjects)* Each process generates the sketches from 𝒮_*local*_, and inserts them into 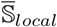, which holds all the sketches generated from that process.

S3) *(gather sketch)* In a global communication step that uses the `MPI_Allgatherv` primitive, we perform a union of all the 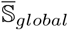 into a single 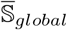 that is stored at each process. Note that each 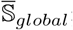 consists of *T* lists, one for each trial, as shown in Fig. 2.

S4) *(map queries)* Each process then processes its local query set 𝒬_*local*_. The mapping stage for each query *q* ∈𝒬_*local*_ comprises of three steps:

Step a) slide window and generate its JEM sketches;

Step b) lookup the subject hits in 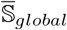 and

Step c) report mapping for the (or a) best hit.

As shown in Fig. 2, hits are located within 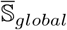 by the corresponding trial numbers (step b). Subsequently, a reporting step scans the bins (or the list of trials) to generate the mapping output pairing queries to subjects (step c).

#### Additional remarks on our parallel implementation

The output for `L2C` mapping is the best hit as shown in Fig. 2. For step (c) above, we implemented a lazy update strategy to support a fast tracking of subjects and their hit rates *across queries*. More specifically, we initialize an array *A*[1, *n*] of tuples of the form ⟨*u, v*⟩, where *u* is an integer counter initialized to 0, and *v* is the query id (initialized to -1). Whenever a query *j* generates a hit with a subject *i*, we check if *A*[*i*].*v* is equal to *j*. If it is, then we simply increment counter *A*[*i*].*u*. But if it is not, then we first set *A*[*i*].*v* to *j*, reset counter *A*[*i*].*u* to 0, and then increment that counter (to 1). This lazy strategy avoids the cost of resetting the counters for all subjects whenever a new query is processed. Note that at each process, queries in 𝒬_*local*_ are processed one by one.

For `L2L`, for a query long-read *l*_*q*_, all the mapped subject reads with a certain frequency ≥ *τ* are reported. The parameter *τ* is the minimum number of trials (out of *T*) during which a query has to generate a hit with a given subject, for it to be considered a successful map. Furthermore, note that for `L2L`, we do not need to process the query set and subject set separately. In the input loading step (S1), we just load the input set of long-reads 𝒮 once, in a distributed manner. Let 𝒮_*local*_ denote the set of local long-reads. When we sketch the subjects (S2), we keep an additional boolean flag, a *tag*, to indicate if the *sketch* has been generated from a *segment* (described in §4.3) or not with each *sketch*. The *gather sketch* step (S3) stays the same. In the *map queries* step (S4), 𝒮_*local*_ is treated as the local query set. The *tag* for each sketch in 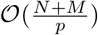 helps to indicate sketches generated from a *segment* of a query long-read *l*_*q*_ (so duplicate work can be avoided). The reporting step scans the bins (or the list of trials) to generate the mapping output pairing long-reads.

### 4.5 Complexity analysis

The runtime complexity analysis of our parallel algorithm is as follows. The input loading step (S1) costs 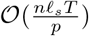 time. Sketching the subjects (S2) can be achieved in 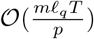 time, where *ℓ*_*s*_ is the average length of a subject. Similarly, sketching the queries (S4) can be achieved in 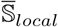 time, where *ℓ*_*q*_ is the average length of a query. The gather step (S3) involves communicating each 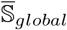 to all processes, and can be achieved in 𝒪 (*λ* log *p* + *μnT*) time, where *λ* is the cost of network latency and *μ* is the reciprocal of network bandwidth (i.e., number of seconds per byte transferred). The parameters *λ* and *μ* are network constants in the Hockney model for parallel performance [38], and they can be empirically determined. While the worst-case size of 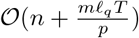 is 𝒪 (*nℓ*_*s*_*T*), in practice we expect significantly fewer minhashes because we are selecting from the list of minimizers *M*_*o*_(*s, w*) (and not all *k*-mers). Finally, the query mapping step (S4) is a local step processing each query *r* ∈𝒬_*local*_. The initialization of counters for the subjects takes 𝒪 (*n*) time and after that each query is mapped through a linear scan of its sequence with *T* minhash computations at all its minimizers. Linear scan is sufficient because of the lazy counter update strategy described above.

Consequently, step S4 takes 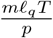 time. For `L2C`, since the number of long reads (*m*) can be expected to be significantly more than the number of contigs (*n*) due to sequencing coverage, we expect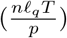 to dominate over *n* in practice. For `L2L`, we are not loading queries or sliding windows over queries seperately as subjects and queries are both long reads. We expect the generating sketches for subjects 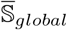 to be the time consuming step for `L2L`.

The space complexity of our approach is dominated by the size of 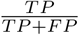. Let *m*_*s*_ denote the average number of minimizers generated per subject. Since we enumerate fixed-size intervals and store one minhash per interval, a subject *s* can be expected to contribute up to 𝒪 (*m*_*s*_*T*) minhashes into the sketch. Therefore, the space complexity per process is 𝒪 (*nm*_*s*_*T*).

## 5 Experimental Results

In this section, we present a thorough experimental evaluation of our sketch-based mapping algorithm, `JEM-mapper` (§ 4.2)). We study both `L2C` and `L2L` use-cases, analyzing the method’s quality and performance and comparing against respective state-of-the-art tools, and using both simulated and real-world data sets.

### 5.1 Experimental setup

#### Test inputs

We used two sets of long read inputs in our experiments (see Table 2):

**Table 2.** Input data sets used in our experiments. All contigs were produced by running Minia assembler [17] on simulated short reads. The long reads are either simulated (default) or real (*O. sativa chr 8 (real)*).

| Input genome |  | S: Subject statistics (Minia contigs) |  |  | Q: Query statistics (HiFi long reads) |  |  |
| --- | --- | --- | --- | --- | --- | --- | --- |
| Input | Genome length (in bp) | No. contigs ($\geq 500$bp) | Total subject size in bp (M) | Contig length (avg.±std.dev) | No. long reads (n) | Total query size in bp (N) | Read length (avg.±std.dev) |
| *E. coli* | 4,641,652 | 375 | 4,510,510 | 12,398 ± 13,010 | 4,010 | 46,005,093 | 10,205 ± 3,418 |
| *P. aeruginosa (unmasked)* | 6,264,404 | 461 | 6,105,989 | 13,782 ± 19,218 | 6,122 | 62,504,041 | 10,221 ± 3,363 |
| *C. elegans* | 100,286,401 | 32,940 | 76,721,376 | 2,330 ± 2,962 | 97,103 | 1,000,011,602 | 10,205 ± 3,400 |
| *D. busckii* | 118,492,362 | 39,352 | 106,278,105 | 2,713 ± 3,174 | 103,781 | 1,058,889,285 | 10,168 ± 3,412 |
| *Human chr 7* | 159,345,973 | 46,798 | 56,547,853 | 1,209 ± 834 | 136,357 | 1,393,462,533 | 9,612 ± 2,988 |
| *Human chr 8* | 145,138,636 | 45,470 | 53,060,920 | 1,167 ± 876 | 141,102 | 1,440,205,836 | 10,200 ± 3,402 |
| *B. splendens (unmasked)* | 339,050,970 | 98,160 | 339,804,114 | 3,462 ± 4,181 | 429,520 | 4,371,221,619 | 10,177 ± 3,403 |
| *O. sativa* | 373,878,990 | 111,788 | 175,228,307 | 1,568 ± 1,446 | 309,615 | 3,012,444,385 | 8,989 ± 62 |
| *Z. mays chr 1* | 307,014,717 | 32,411 | 50,255,597 | 1,550 ± 1,332 | 301,607 | 2,703,145,070 | 8,995 ± 35 |
| *O. sativa chr 8 (real)* | 28,443,022 | 9,945 | 18,416,389 | 1,851 ± 2,067 | 532,667 | 10,458,872,536 | 19,642 ± 4,246 |

- *PacBio HiFi simulated long reads:* These are read generated using the PBSIM3 PacBio read simulator [39]. PBSIM3 generates SAM format data for CLR reads, which is then converted to BAM files and, finally using bioconda [40] pacakge *pbccs*, CCS (Circular Consensus Sequencing) HiFi reads are generated. Simulations were run with a low coverage of 10× and a long read median length 10Kbp; and
- *PacBio HiFi real long reads:* These is a collection of real-world PacBio HiFi reads for *Oryza sativa* (chr 8), downloaded from the PacBio repository [41].

The simulated read data sets allow us to evaluate using a ground-truth (using the coordinate information from SAM file), while the real-world data set is aimed at a real-world application. Simulated reads were generated from real-world whole genomes, downloaded from NCBI GenBank [42], for eight different organisms ranging from bacterial to eukaryotic species (listed in Table 2). For a subset of these genomes (with the exception of *P. aeurginosa* and *B. splendens*), repeat information was available, and so we masked those inputs using the RepeatMasker tool [36]. For the *P. aeruginosa* and *B. splendens* genomes, since repeat information was not available, we used unmasked inputs.

We used the following two steps to construct the contigs for all the inputs: use the ART sequencing simulator [43] to generate 100bp Illumina short reads; and assemble the short reads using the Minia assembler [17] into contigs.

#### Test platform

All experiments were conducted on a distributed memory cluster with 9 compute nodes, each with 64 AMD Opteron™ (2.3GHz) cores and 128 GB DRAM. The nodes are interconnected using 10Gbps Ethernet and share 190TB of ZFS storage. The cluster supports OpenMPI (for distributed memory MPI codes) and OpenMP (for shared memory multithreaded codes).

#### Software configuration

All runs of our software `JEM-mapper` was performed using the following set of parameters as default: *k* = 16 bp; no. trials *T* = 30 (choice explained in Fig. 6); and a window size of *w* = 100 bp to generate minimizer sketches. In other words, we select a *k*-mer (of size 16 bp) from a consecutive stretch of *w* (100) *k*-mers to be the minimizer (as explained in Section 4.2). These minimizers are then added to the corresponding set *M*_*o*_(*s, w*) only if they change or if the current minimizer goes out of bounds. Subsequently, to generate the JEM sketches (Algorithm 1), we set the interval length same as the segment *ℓ* bp (for `L2C` 1,000 bp and for `L2L` 2,000 bp) for long reads. The smaller *ℓ* length for `L2C` is to help capture overlaps with shorter contigs. Setting *ℓ* = 2, 000 bp for the `L2L` is consistent with other long read mappers [15, 18]. For `L2L`, the minimum number of trials needed to generate a hit (i.e., *τ*) was set to 15 (out of *T* = 30 trials), to represent a 50% hit rate with the random trials.

### 5.2 Evaluation for L2C mapping

Once the long reads are generated, we pulled out the two end segments (prefix and suffix) of length *ℓ* = 1, 000 bp and added them to the query set 𝒬. For comparative evaluation, the two state-of-the-art reference genome mappers that support `L2C` (see Table 1) are `Mashmap` [21] and `Minimap2` [15]. Of these two, `Mashmap` tool [21] is a fast reference genome mapper that also uses sketching, and from its implementation we can easily extract the top hit for a query, making it possible to directly compare it with `JEM-mapper`. As for `Minimap2` [15], it follows a more classical seed-and-extend alignment-based approach, but it also benefits from the use of minimizers internally for the seeding step. However, it was not possible to make a direct comparison with its output because it reports multiple hits for each query. Therefore, we focus our comparative evaluation on `Mashmap`. In addition to `Mashmap`, we also implemented the classical `MinHash` scheme (Section 4.1) by modifying `JEM-mapper` implementation. This allowed us to compare the efficacy of the Jaccard Estimator MinHash scheme over the classical `MinHash` scheme. In all cases, the *same* inputs (𝒬, 𝒮) were provided to both programs—i.e., mapping the end segments of long reads to contigs.

#### Evaluation methodology

For quality evaluation, we constructed a *benchmark* for all simulated data sets using the coordinate information of the contigs (𝒮) and long reads (𝒬) mapped back to the full-length reference genome (𝒢). This is illustrated in Fig. 4. More specifically, to determine the ⟨start,end⟩ coordinates of each contig, we mapped the set of contigs to the reference 𝒢 using Minimap2 [15]. The coordinates of the long reads are readily available from SAM files (generated by PBSIM3). A given end segment of a long read *e* ∈ 𝒬 is said to *map* to a contig *c* ∈ 𝒮 if and only if its respective ⟨start,end⟩ coordinates overlap over at least *k* base pairs of the reference genome—as shown in Fig. 4.

**Figure 4.**
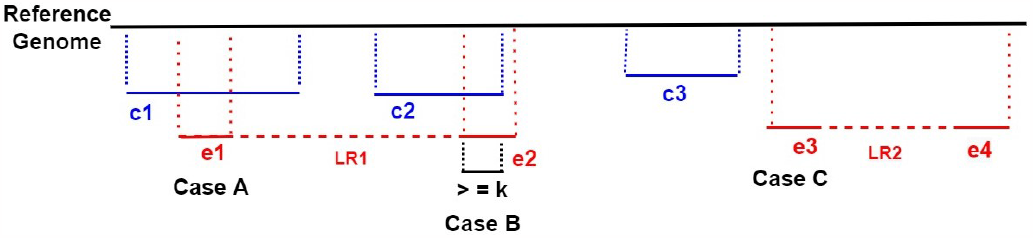
`L2C`: *Benchmark* cases for when an ending segment of a long read is said to successfully map (Cases A and B) or *not* map (Case C) to a contig. In the figure, two long reads are shown, one with end segments ⟨*e*1, *e*2⟩ and another with end segments ⟨*e*3, *e*4⟩.

Let Bench denote the set of all true ⟨read end,contig⟩ mappings. Let Test denote the set of output ⟨read end,contig⟩ mappings produced by one of the test implementations. Then, we classify each ⟨read end *e*,contig *c*⟩ pair as:

- **True Positive (TP):** if ⟨*e, c*⟩ ∈ Test and ⟨*e, c*⟩ ∈ Bench
- **False Positive (FP):** if ⟨*e, c*⟩ ∈ Test and ⟨*e, c*⟩ ∉ Bench
- **False Negative (FN):** if ⟨*e, c*⟩ ∉ Test and ⟨*e, c*⟩ ∈ Bench
- **True Negative (TN):** if ⟨*e, c*⟩ ∉ Test and ⟨*e, c*⟩ ∉Bench

Based on the above four measures, we calculate *precision* as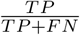 and *recall* as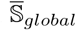. Note that if an output mapping is a false positive, then by implication it is also a false negative (since there is room for only one best hit in the `L2C` case). But there can be additional false negatives. Therefore the recall values are upper bounded by the precision values in this evaluation scheme.

#### Qualitative evaluation

Fig. 5 shows the precision (left) and recall (right) values for `JEM-mapper` and `Mashmap`. These results are for the PacBio HiFi simulated long reads. The results show that by and large, our sketch-based implementation is competitive and show comparable quality compared to `Mashmap` in all cases, with both tools producing well over 98% precision for all inputs tested. For the smaller/less complex genomes (*E. coli, P. aeruginosa (unmasked)*) `JEM-mapper` produces similar precision values as `Mashmap`. Our scheme produces slightly better precision for all the larger (more complex) inputs. As mentioned earlier, for *B. splendens*, we have used unmasked input and we can see from the Fig. 5 (left), that `JEM-mapper` is outperforming `Mashmap`.

**Figure 5.**
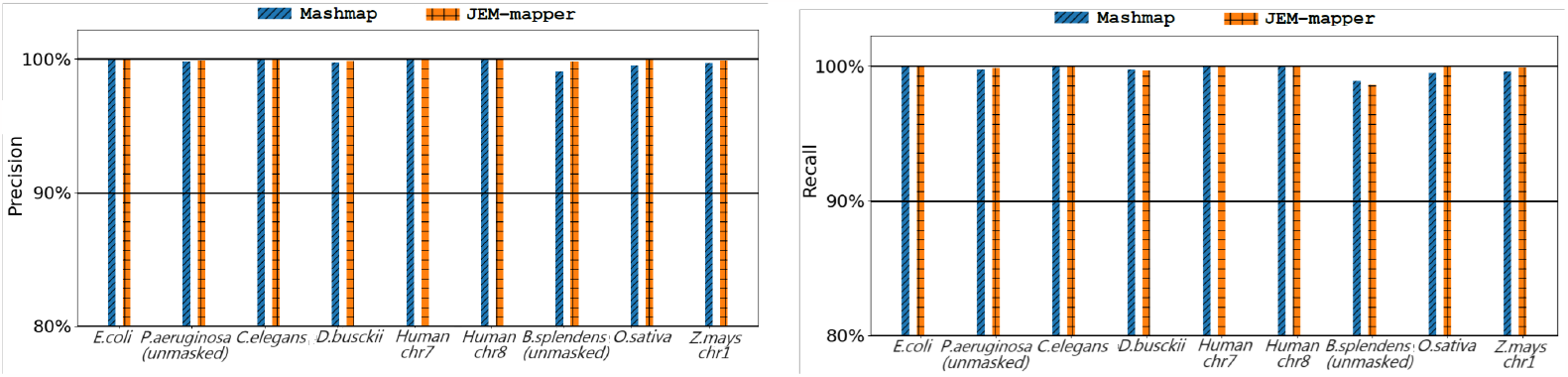
`L2C`: Comparative quality evaluation (precision and recall), comparing `JEM-mapper` and `Mashmap`, on various PacBio HiFi simulated long reads inputs.

**Figure 6.**
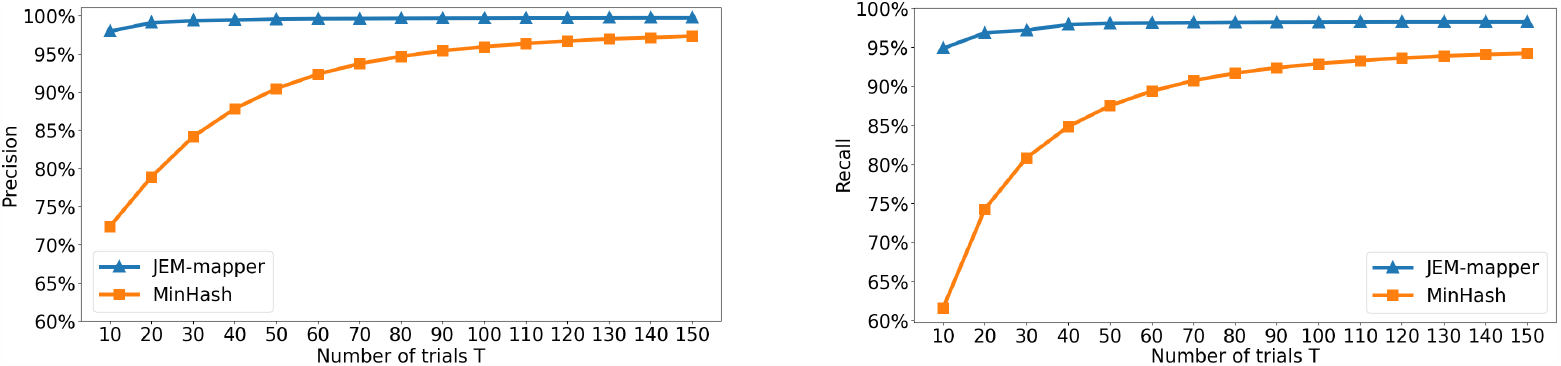
`L2C`: Effect of varying the number of trials on quality results (precision and recall), using input *B. splendens (unmasked)*. We can observe that `JEM-mapper` can achieve above 97% precision and recall only using 20 trials whereas classical `MinHash` needed to use more than 150 trials to reach a similar quality output.

Eukaryotic inputs have more repetitive content that may lead to reduced precision and the results show that the strategy to select the best candidate from multiple random trials makes our sketch-based scheme more precise for these more complex inputs.

For all the input datasets, `JEM-mapper` produces similar recall as compared to `Mashmap`. The difference between the two tools is marginal in all cases (except for *B. splendens (unmasked)*). Again, both tools produce recall values that are 98% or more for most inputs. We also note that the recall values are very close to the precision values, implying that most of the loss in recall can be attributed to false positive mapping in the top hit. Note that if we are to extend our method to report a fixed number, say top *x* hits per read, then several of the missing contig hits could possibly be recovered and recall improved.

The number of trials *T* could have an impact on the overall quality. Fig. 6 shows the effect of varying *T* on the `JEM-mapper` and classical `MinHash` implementation, using the *B. splendens (unmasked)* input. Increasing *T* improves both precision and recall. This behavior is consistent with the fact that with more trials, the sketch-based schemes get more chances to find a hit between a long read and a contig, thus making the recall better. Since these are 99.9% correct long reads, precision also improves. We can observe that `JEM-mapper` can achieve above 97% precision and recall only using 20 trials. After 30 trials it reaches to saturation for precision and recall values and adding more trials only improves precision and recall marginally. However, for classical `MinHash`, even after using 100 iterations, the precision and recall values are quite low compared to `JEM-mapper`—demonstrating the advantage of the Jaccard Estimator Minhash scheme over classical `MinHash`. This behavior is expected as `JEM-mapper` is better equipped to identify the region within *ℓ*-long segment stretches of the subject. In contrast, classical `MinHash` does not constrain identification of sketches from within such distance bounds, and therefore, may need more random trials to recover the hits. The property of supporting fewer number of trials also provides a significant advantage to `JEM-mapper` toward faster performance. For example, to achieve roughly the same quality in mapping on the *B. splendens (unmasked)* input, `JEM-mapper` took 30 trials, whereas the classical `MinHash` implementation took 150 trials.

#### Performance evaluation

Next, we evaluate the runtime and parallel performance of `JEM-mapper` and compare it with state-of-the-art tools. First, we studied the strong scaling behavior of our parallel implementation for `JEM-mapper`, by varying *p* from 4 through 64. Table 3 shows the parallel runtimes for the larger inputs tested. Overall, the parallel runtime reduces with increase in *p*, demonstrating improving speedups. For instance, on *B. splendens (unmasked)*, the relative speedup (relative to *p* = 4) increases from 1.9× on *p* = 8, to 2.9× on *p* = 16, 3.5× on *p* = 32, and 4.1× on *p* = 64. As the number of processes increases, the work per process also reduces leading to parallel overheads slowly starting to dominate. We have compared our distributed memory implementation results with `Mashmap` runtimes. `Mashmap` only supports shared memory parallelism using multithreading. Table 3 shows the `Mashmap` runtimes for where the number of threads is set to 64. The results show that `JEM-mapper` is significantly faster than the `Mashmap` implementations. In all the input cases, `JEM-mapper` running in distributed memory mode with *p* = 64 yields higher speedup (ranging from 5.6× to 13×) over `MashMap` running on the same number of processors (no. threads = 64). Note that in parallel processing, distributed memory setting is expected to have more overheads due to network communication.

**Table 3.** `L2C`: Strong scaling results for `JEM-mapper`: Shown are the parallel runtimes (in sec) for `JEM-mapper` as function of the number of processes (*p*) on various inputs. Also shown are the `Mashmap` runtimes, which was run on 64 threads (*t*), as the tool supports only shared memory parallelism.

| Input | JEM-mapper |  |  |  |  | Mashmap |
| --- | --- | --- | --- | --- | --- | --- |
| | $p = 4$ | $p = 8$ | $p = 16$ | $p = 32$ | $p = 64$ | $t = 64$ |
| *C. elegans* | 125 | 54 | 41 | 27 | 24 | 227 |
| *D. busckii* | 155 | 85 | 57 | 36 | 33 | 251 |
| *Human chr 7* | 128 | 75 | 39 | 27 | 23 | 260 |
| *Human chr 8* | 119 | 63 | 39 | 25 | 22 | 257 |
| *B. splendens (unmasked)* | 519 | 280 | 180 | 145 | 127 | 876 |
| *O. sativa* | 557 | 216 | 130 | 83 | 69 | 677 |
| *Z. mays chr 1* | 283 | 164 | 99 | 54 | 44 | 528 |
| *O. sativa chr 8 (real)* | 420 | 218 | 122 | 91 | 69 | 605 |

Fig. 7a (left) shows the parallel runtime broken down by the individual steps of the `JEM-mapper` implementation for *p* = 16. It is evident that the dominant step is the query processing time, which includes the time to sketch the queries and search in the hash table and report the hits.

**Figure 7.**
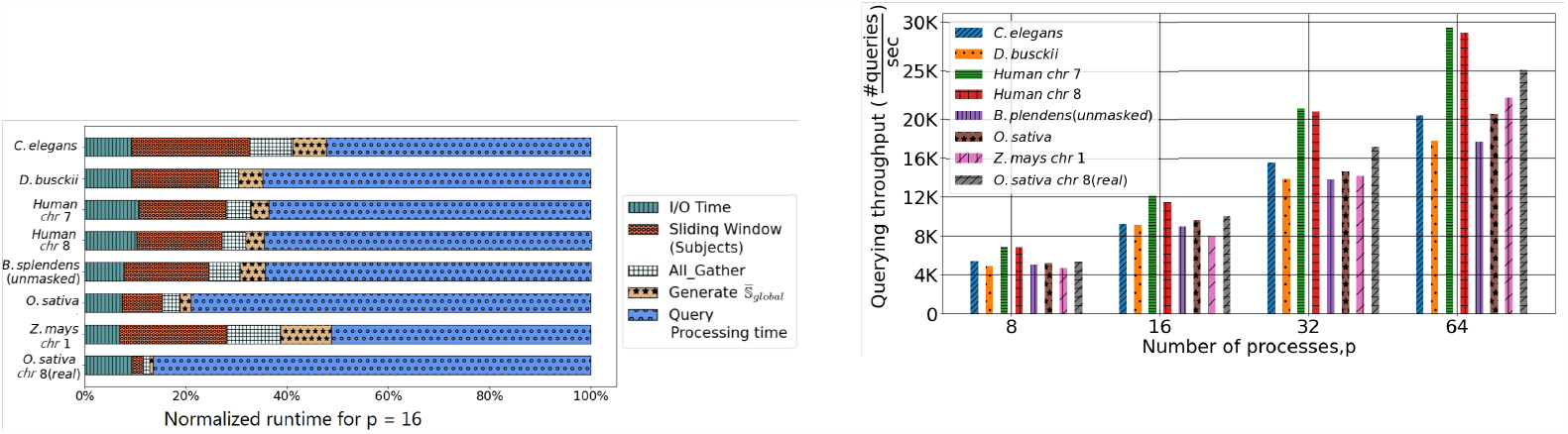
`L2C`: (a) Runtime breakdown by steps for `JEM-mapper`; (b) Querying through-put for `JEM-mapper`.

We also closely analyzed the query processing time from the perspective of querying throughput, defined as the number of queries processed per unit time (sec). To calculate this, we included the times for sliding windows on the queries, sketching the queries, and search in 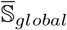 and report step. Fig. 7b (right) shows the querying throughput for our `JEM-mapper` implementation, for the larger inputs tested. We observe that this querying throughput scales almost linearly.

Fig. 8 shows the total computation versus communication time for *Human chr 7* and *B. splendens (unmasked)* varying the number of processors from *p* = 4 to *p* = 64. The total computation time includes the I/O time, subject processing time, generating the 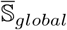 time, and the query processing and search time. As expected, increasing the number of processors increases the total communication overhead, but the overhead stays well under 25% for up to *p* = 64.

**Figure 8.**
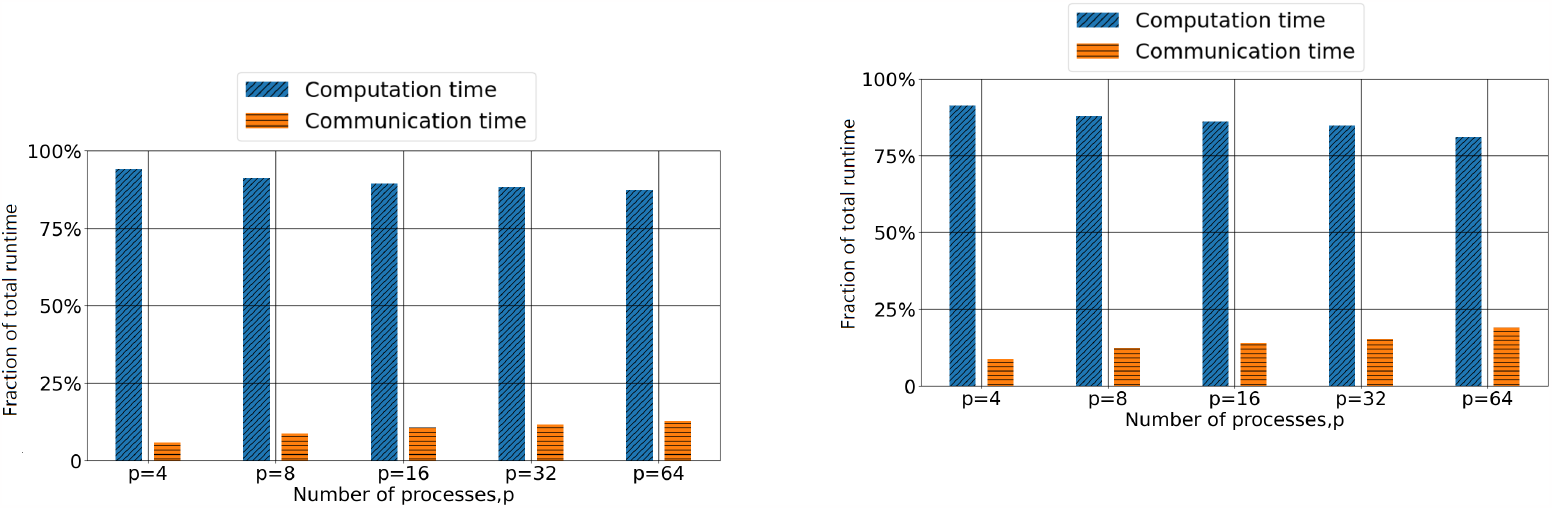
`L2C`: The fractions of the total runtime spent in computation (blue) versus communication (orange) for (a) *Human chr 7*; and (b) *B. splendens (unmasked)*. For a scalable parallel execution, computation time should dominate over communication.

#### Evaluation on real-world data

As a real-world application, we used a real-world PacBio HiFi long read data set for *Oryza sativa* (rice MH63), downloaded from the PacBio repository [41] (see Table 2 for input statistics). We used only the long reads from chromosome 8 for our experimental analysis (*n* = 532*K*). We then used `JEM-mapper` to map these long reads to the contigs generated using a Minia assembly of *O. sativa* chr. 8 short reads (*m* = 9.9*K*). Finally, we used BLAST [44] to compute the percent identity between each long read (segment) and the corresponding mapped contig as reported by `JEM-mapper`. Fig. 9 shows that the percent identity between most of the long read ends and the corresponding contig falls between 95%-100%—showing the high quality of mapping generated by `JEM-mapper`.

**Figure 9.**
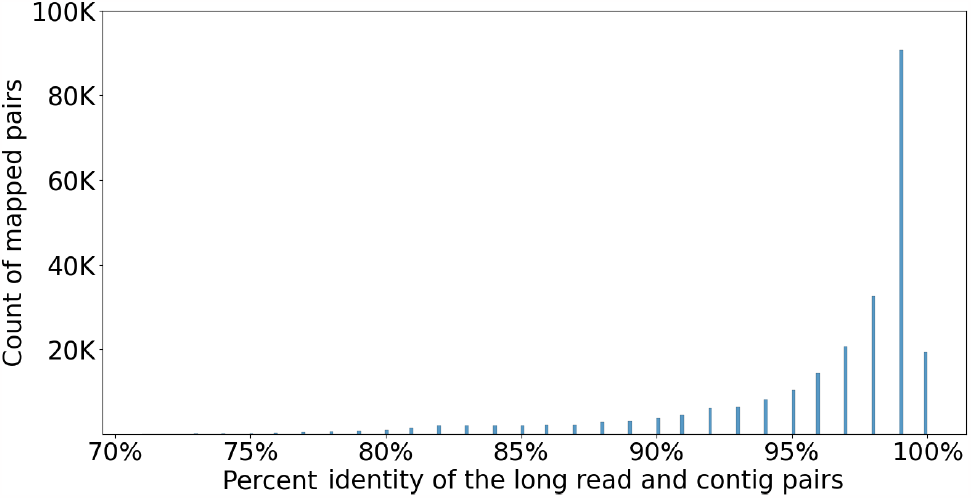
`L2C`: Percent identity distribution for long read mappings generated by `JEM-mapper` on the *O. sativa* data set.

### 5.3 Evaluation for L2L mapping

In the `L2L` use-case, the input consists of only a set of long reads. We have used the same synthetic and real-world data as mentioned in (Section 5.1). For comparative evaluation, we compared against two state-of-the-art long read overlap detection tools, namely `Minimap2` [15] and `MECAT` [18]—as noted in Table 1. As mentioned earlier for `Minimap2` [15], it follows a classical seed and extend-based approach, but it also benefits from the use of minimizers internally for the seeding step. `MECAT` [18] is an alignment-free approach that relies on *k*-mers to detect overlapping candidates and uses a pseudo-linear alignment scoring algorithm to discard false overlap candidates. In all cases, the *same* inputs (𝒮) were provided to all the tools.

#### Evaluation methodology

For quality evaluation in the `L2L` use-case, we constructed a *benchmark* using the genomic coordinate information of the long reads. First, all the input long reads (𝒮) were positioned, using the coordinate information from PBSIM3, along the reference genome 𝒢. A query long read *l*_*q*_ is said to *map* to a long read *l* ∈ 𝒮 if and only if their respective <start,end> coordinates overlap in at least 2 Kbp positions of the reference genome—as shown in Fig. 10. This overlap cutoff is derived from the state-of-the-art tools [8, 37].

**Figure 10.**
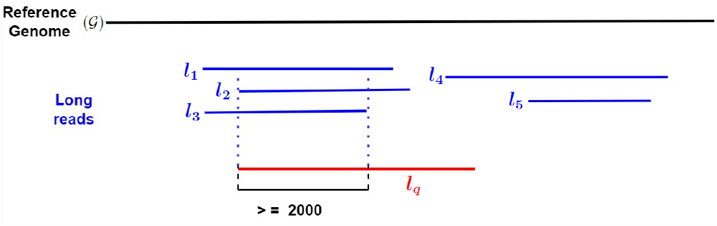
`L2L`: In our *benchmark* each query long read *l*_*q*_ is said to map to a subset of long reads in 𝒮 ({*l*_1_, *l*_2_, *l*_3_}in the example) if the overlap is more than 2 Kbp. Note that, based on this criterion, *l*_*q*_ does *not* map to *l*_4_ and *l*_5_.

Let Bench denote the set of all true mappings generated as above. Let Test denote the set of all test mappings generated by our implementation. Then, we can place each distinct <*l*_*q*_, *l*> pair into one of the following categories: TP, FP, FN, and TN, and subsequently calculate precision and recall—consistent with our definitions in Section 5.2. Note that since this `L2L` allows each query long read to map to one or more long reads, a false positive does not necessarily imply a false negative (as it did for `L2C`).

#### Qualitative evaluation

Fig. 11 shows the precision (left) and recall (right) values for `JEM-mapper`, with comparisons to `Minimap2` and `MECAT`. These results are for the PacBio HiFi simulated long reads. The results indicate that, for the most part, `JEM-mapper` exhibits comparable quality to whichever tool is the best between `Minimap2` and `MECAT` for each input. From Fig. 11 (left) we observe that all precision values are generally well above 88% for masked inputs using `JEM-mapper`. For all the input data sets, the recall values are between 85% to 99.9% using `JEM-mapper`. For the smaller/less complex genomes (*E. coli, P. aeruginosa (unmasked)*), `JEM-mapper` produces similar recall values as `MECAT`.

**Figure 11.**
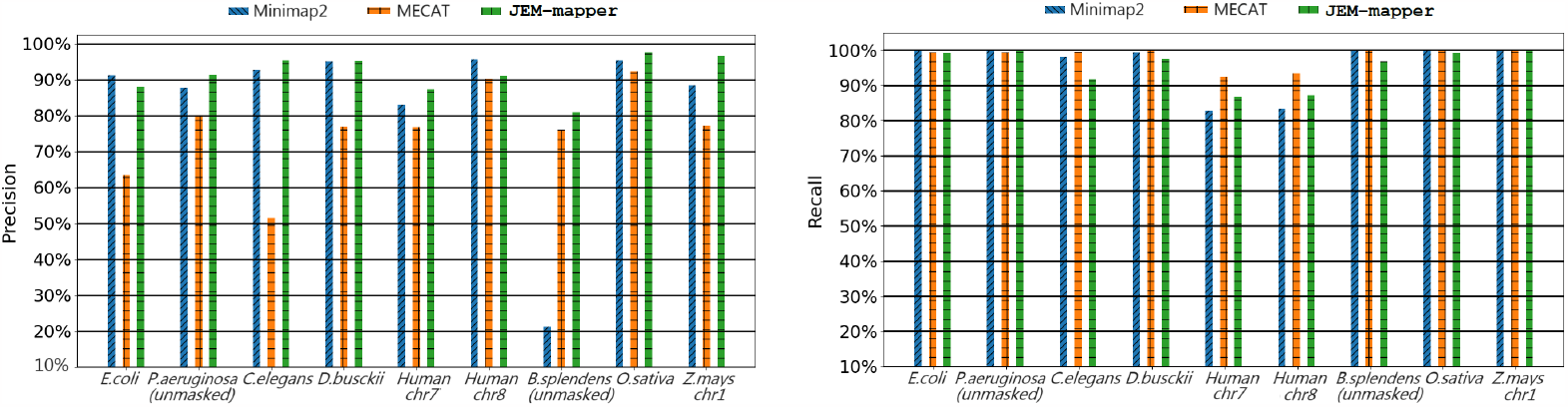
`L2L`: Quality results (precision and recall) using PacBio HiFi simulated long reads for different long-read overlappers

As for recall, Fig. 11 (right) shows that all three tools perform comparably, with `MECAT` yielding better recall values for some of the inputs. However, its precision is also lower than the other two tools for many of the inputs. In contrast, `JEM-mapper` and `Minimap2` perform consistently (precision and recall-wise) for the masked inputs. Note that for unmasked input *B. splendens (unmasked)*, `Minimap2` attain a very low precision (near 20%) since the repetitive regions of the genome can possibly produce many false-positive overlaps. `JEM-mapper` overcomes this issue by using the frequency-based heuristic (described in (Section 4.3)).

#### Performance Evaluation

We studied the strong scaling behavior of our parallel implementation for `JEM-mapper`, by varying *p* from 4 through 64. Table 4 shows the parallel runtimes for the larger inputs tested. As with our `L2C` study, the results show good parallel scaling behavior for `JEM-mapper`. As a concrete example, for the *O. sativa* input, the relative speedup (relative to *p* = 4) increases from 1.9× on *p* = 8 to 6.3× on *p* = 64.

**Table 4.** `L2L`: Strong scaling results for `JEM-mapper`: Shown are the parallel runtimes (in sec) for `JEM-mapper` as function of the number of processes (*p*) on various inputs. Also `Minimap2` and `MECAT` runtimes are shown. These tools were run on 64 threads (*t*), as the tool supports only shared memory parallelism.

| Input | JEM-mapper |  |  |  |  | Minimap2 | MECAT |
| --- | --- | --- | --- | --- | --- | --- | --- |
| | $p = 4$ | $p = 8$ | $p = 16$ | $p = 32$ | $p = 64$ | $t = 64$ | $t = 64$ |
| *C. elegans* | 485 | 264 | 154 | 101 | 79 | 73 | 498 |
| *D. busckii* | 589 | 318 | 218 | 160 | 102 | 74 | 595 |
| *Human chr 7* | 605 | 323 | 205 | 137 | 115 | 60 | 470 |
| *Human chr 8* | 707 | 295 | 178 | 119 | 103 | 71 | 439 |
| *B. splendens (unmasked)* | 2,521 | 1,378 | 760 | 494 | 388 | 2,886 | 2,931 |
| *O. sativa* | 1,902 | 1,030 | 579 | 376 | 302 | 3,008 | 2,184 |
| *Z. mays chr 1* | 1,894 | 945 | 528 | 350 | 281 | 4,123 | 2,670 |
| *O. sativa chr 8 (real)* | 3,171 | 1,659 | 947 | 608 | 494 | 7,479 | 3,418 |

Table 4 also compares the parallel runtimes achieved by `JEM-mapper` to the corresponding multithreaded parallel runtimes achieved by `Minimap2` and `MECAT`. For the smaller inputs, `Minimap2` shows faster runtimes than `JEM-mapper`. However, for the larger inputs, it is evident that the runtimes for `Minimap2` significantly increases.

Whereas `JEM-mapper` outperforms both `Minimap2` and `MECAT` on these larger inputs. For instance, for *O. sativa*, `JEM-mapper` running on distributed memory with *p* = 64 yields 9.9× and 7.2× speedups over `Minimap2` and `MECAT` running on 64 threads, respectively.

Furthermore, comparing Tables 3 and 4, we observe that the `L2L` runtimes are larger than the `L2C` runtimes for the corresponding inputs. This is to be expected because the entire long read set is treated as both the subject set and query set for `L2L`, implying more work.

Fig. 12 shows the parallel runtime broken down by the individual steps of the `JEM-mapper` implementation for *p* = 16. It is evident that the dominant step is the sliding window of the subjects, which includes the time to sketch the subjects. Query processing time includes the search in 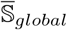and the report step. Note that the fraction of total runtime spent in query processing is significantly smaller compared to the corresponding fractions observed in the `L2C` case (Figure 7 (left)). This is because our parallel implementation is optimized to avoid duplicated effort in the sketching of query processing, since 𝒮 = 𝒬 for `L2L` (as noted in Section 4.4).

**Figure 12.**
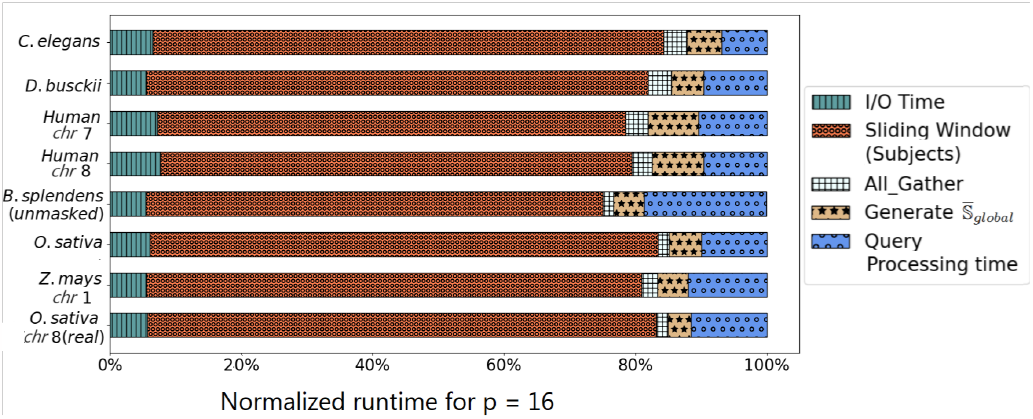
`L2L`: Normalized runtime breakdown by steps for `JEM-mapper` implementation for *p* = 16

Fig. 13 shows the total computation versus communication time for *O. sativa* and *Z. mays chr 1* varying the number of processors from *p* = 4 to *p* = 64. The total computation time includes the I/O time, subject processing time, generating the 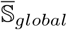 time, and the query processing time. As we have seen earlier, increasing the number of processors increases the total communication overhead, but the overhead stays well under 10% for even up to *p* = 64.

**Figure 13.**
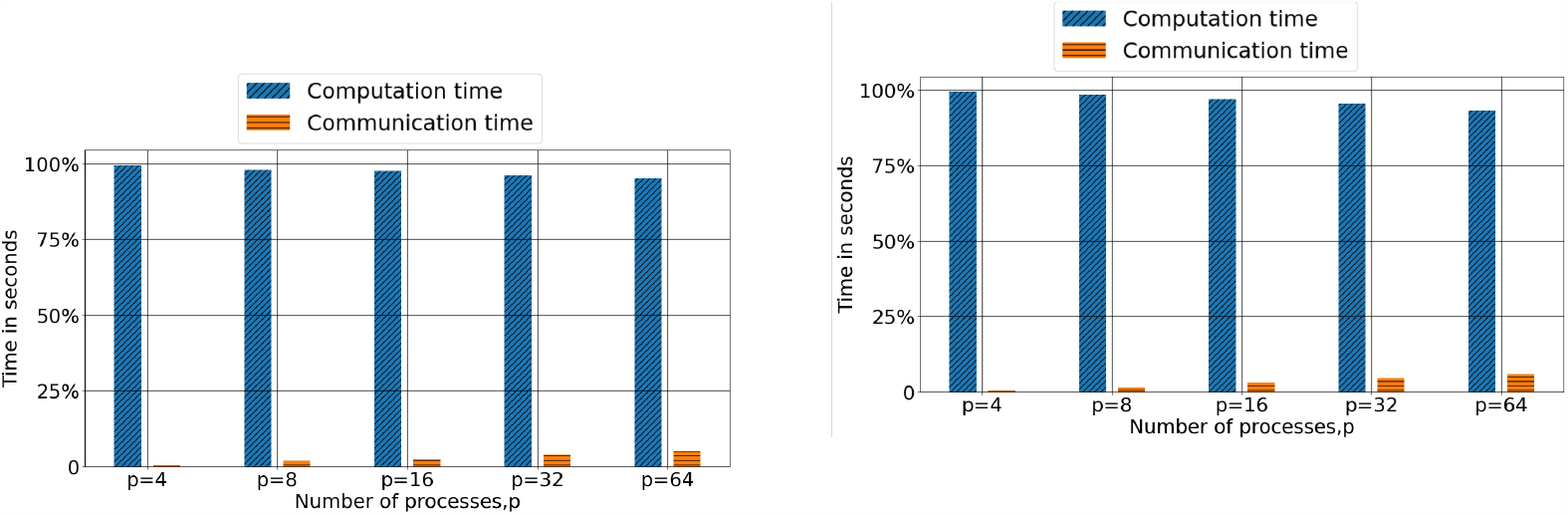
`L2L`: The fractions of the total runtime spent in computation (blue) versus communication (orange) for (a) *O. sativa*; and (b) *Z. mays chr 1*;

## 6 Conclusions

In this paper, we presented a minimizer-based Jaccard estimator sketch-based algorithm for mapping long reads to different types of biological sequences. Our mapping algorithm can be used to map long reads to either a set of partially assembled contigs (from a previous short read assembly), or to the set of long reads themselves. The former application can be used to extend the reach of previously constructed short read assemblies (using long reads), while the latter can be used in *de novo* long read assembly workflows. All our results indicate that we are able to meet the quality in both precision and recall, compared to the corresponding state-of-the-art mapping tools, while providing significant advantages in performance speedup. Our open source parallel implementations can be executed on distributed memory platforms (i.e., clusters) which make them suited to scale to large inputs.

This work has opened up multiple avenues for future research, including (but not limited to): i) integration into end-to-end hybrid assembly and scaffolding workflows; and ii) large-scale studies targeting more complex eurakyotic genomes; iii) extension to reference-guided assembly pipelines [13], where either reads are mapped against the reference genome or alternatively contigs or scaffolds are aligned against the reference genome. These use-cases can further enhance the utility of this standalone mapping tool, and help in better harnessing the power of long read sequencing into existing assembly and sequencing workflows.

## Acknowledgments

This research was supported in parts by NSF grants OAC 1910213 and CCF 1919122. We thank Dr. Priyanka Ghosh for several discussions during the early stages of the project.

## References

1. C. E. Mason and O. Elemento, “Faster sequencers, larger datasets, new challenges,” 2012.

2. D. Deamer, M. Akeson, and D. Branton, “Three decades of nanopore sequencing,” Nature biotechnology, vol. 34, no. 5, pp. 518–524, 2016.

3. M. O. Pollard, D. Gurdasani, A. J. Mentzer, T. Porter, and M. S. Sandhu, “Long reads: their purpose and place,” Human molecular genetics, vol. 27, no. R2, pp. R234–R241, 2018.

4. T. Hon, K. Mars, G. Young, Y.-C. Tsai, J. W. Karalius, J. M. Landolin, N. Maurer, D. Kudrna, M. A. Hardigan, C. C. Steiner, et al., “Highly accurate long-read hifi sequencing data for five complex genomes,” Scientific data, vol. 7, no. 1, pp. 1–11, 2020.

5. P. Morisse, T. LeCroq, and A. LeFeBVre, “Long-read error correction: a survey and qualitative comparison,” BioRxiv, pp. 2020–03, 2021.

6. S. Koren, B. P. Walenz, K. Berlin, J. R. Miller, N. H. Bergman, and A. M. Phillippy, “Canu: scalable and accurate long-read assembly via adaptive k-mer weighting and repeat separation,” Genome research, vol. 27, no. 5, pp. 722–736, 2017.

7. G. M. Kamath, I. Shomorony, F. Xia, T. A. Courtade, and N. T. David, “HINGE: long-read assembly achieves optimal repeat resolution,” Genome research, vol. 27, no. 5, pp. 747–756, 2017.

8. G. Guidi, M. Ellis, D. Rokhsar, K. Yelick, and A. Buluç, “BELLA: Berkeley efficient long-read to long-read aligner and overlapper,” in SIAM Conference on Applied and Computational Discrete Algorithms (ACDA21), pp. 123–134, SIAM, 2021.

9. K. Berlin, S. Koren, C.-S. Chin, J. P. Drake, J. M. Landolin, and A. M. Phillippy, “Assembling large genomes with single-molecule sequencing and locality-sensitive hashing,” Nature biotechnology, vol. 33, no. 6, pp. 623–630, 2015.

10. D. Antipov, A. Korobeynikov, J. S. McLean, and P. A. Pevzner, “hybridSPAdes: an algorithm for hybrid assembly of short and long reads,” Bioinformatics, vol. 32, no. 7, pp. 1009–1015, 2016.

11. E. Haghshenas, H. Asghari, J. Stoye, C. Chauve, and F. Hach, “Haslr: Fast hybrid assembly of long reads,” Iscience, vol. 23, no. 8, p. 101389, 2020.

12. T. Rahman, O. Bhowmik, and A. Kalyanaraman, “An Efficient Parallel Sketch-based Algorithm for Mapping Long Reads to Contigs,” in 2023 IEEE International Parallel and Distributed Processing Symposium Workshops (IPDPSW), pp. 157–166, IEEE, 2023.

13. H. E. Lischer and K. K. Shimizu, “Reference-guided de novo assembly approach improves genome reconstruction for related species,” BMC bioinformatics, vol. 18, no. 1, pp. 1–12, 2017.

14. B. Langmead, C. Trapnell, M. Pop, and S. L. Salzberg, “Ultrafast and memory-efficient alignment of short dna sequences to the human genome,” Genome biology, vol. 10, no. 3, pp. 1–10, 2009.

15. H. Li, “Minimap2: pairwise alignment for nucleotide sequences,” Bioinformatics, vol. 34, no. 18, pp. 3094–3100, 2018.

16. A. V. Zimin and S. L. Salzberg, “The SAMBA tool uses long reads to improve the contiguity of genome assemblies,” PLoS computational biology, vol. 18, no. 2, p. e1009860, 2022.

17. R. Chikhi and G. Rizk, “Space-efficient and exact de bruijn graph representation based on a bloom filter,” Algorithms for Molecular Biology, vol. 8, no. 1, pp. 1–9, 2013.

18. C.-L. Xiao, Y. Chen, S.-Q. Xie, K.-N. Chen, Y. Wang, Y. Han, F. Luo, and Z. Xie, “MECAT: fast mapping, error correction, and de novo assembly for single-molecule sequencing reads,” Nature methods, vol. 14, no. 11, pp. 1072–1074, 2017.

19. C. Jain, S. Koren, A. Dilthey, A. M. Phillippy, and S. Aluru, “A fast adaptive algorithm for computing whole-genome homology maps,” Bioinformatics, vol. 34, no. 17, pp. i748–i756, 2018.

20. G. Myers, “Efficient local alignment discovery amongst noisy long reads,” in International Workshop on Algorithms in Bioinformatics, pp. 52–67, Springer, 2014.

21. C. Jain, A. Dilthey, S. Koren, S. Aluru, and A. M. Phillippy, “A fast approximate algorithm for mapping long reads to large reference databases,” in International Conference on Research in Computational Molecular Biology, pp. 66–81, Springer, 2017.

22. B. D. Ondov, T. J. Treangen, P. Melsted, A. B. Mallonee, N. H. Bergman, S. Koren, and A. M. Phillippy, “Mash: fast genome and metagenome distance estimation using minhash,” Genome biology, vol. 17, no. 1, pp. 1–14, 2016.

23. B. D. Ondov, G. J. Starrett, A. Sappington, A. Kostic, S. Koren, C. B. Buck, and A. M. Phillippy, “Mash screen: high-throughput sequence containment estimation for genome discovery,” Genome biology, vol. 20, no. 1, pp. 1–13, 2019.

24. G. Marçais, D. DeBlasio, P. Pandey, and C. Kingsford, “Locality-sensitive hashing for the edit distance,” Bioinformatics, vol. 35, no. 14, pp. i127–i135, 2019.

25. L. Coombe, J. X. Li, T. Lo, J. Wong, V. Nikolic, R. L. Warren, and I. Birol, “LongStitch: High-quality genome assembly correction and scaffolding using long reads,” BMC bioinformatics, vol. 22, pp. 1–13, 2021.

26. M. Roberts, W. Hayes, B. R. Hunt, S. M. Mount, and J. A. Yorke, “Reducing storage requirements for biological sequence comparison,” Bioinformatics, vol. 20, no. 18, pp. 3363–3369, 2004.

27. A. Z. Broder, “On the resemblance and containment of documents,” in Proceedings. Compression and Complexity of SEQUENCES 1997 (Cat. No. 97TB100171), pp. 21–29, IEEE, 1997.

28. G. Holley, R. Wittler, J. Stoye, and F. Hach, “Dynamic alignment-free and reference-free read compression,” Journal of Computational Biology, vol. 25, no. 7, pp. 825–836, 2018.

29. G. Marçais, D. Pellow, D. Bork, Y. Orenstein, R. Shamir, and C. Kingsford, “Improving the performance of minimizers and winnowing schemes,” Bioinformatics, vol. 33, no. 14, pp. i110–i117, 2017.

30. M. Belbasi, A. Blanca, R. S. Harris, D. Koslicki, and P. Medvedev, “The minimizer jaccard estimator is biased and inconsistent,” bioRxiv, 2022.

31. M. Cechova, “Probably correct: rescuing repeats with short and long reads,” Genes, vol. 12, no. 1, p. 48, 2020.

32. M. J. Chaisson and G. Tesler, “Mapping single molecule sequencing reads using basic local alignment with successive refinement (BLASR): application and theory,” BMC bioinformatics, vol. 13, no. 1, pp. 1–18, 2012.

33. R. Guo, Y.-R. Li, S. He, L. Ou-Yang, Y. Sun, and Z. Zhu, “Replong: de novo repeat identification using long read sequencing data,” Bioinformatics, vol. 34, no. 7, pp. 1099–1107, 2018.

34. P. S. Schnable, D. Ware, R. S. Fulton, J. C. Stein, F. Wei, S. Pasternak, C. Liang, J. Zhang, L. Fulton, T. A. Graves, et al., “The B73 maize genome: complexity, diversity, and dynamics,” Science, vol. 326, no. 5956, pp. 1112–1115, 2009.

35. B. Sosinski, V. Shulaev, A. Dhingra, A. Kalyanaraman, R. Bumgarner, D. Rokhsar, I. Verde, R. Velasco, and A. G. Abbott, “Rosaceaous genome sequencing: perspectives and progress,” Genetics and genomics of Rosaceae, pp. 601–615, 2009.

36. N. Chen, “Using repeat masker to identify repetitive elements in genomic sequences,” Current protocols in bioinformatics, vol. 5, no. 1, pp. 4–10, 2004.

37. H. Li, “Minimap and miniasm: fast mapping and de novo assembly for noisy long sequences,” Bioinformatics, vol. 32, no. 14, pp. 2103–2110, 2016.

38. R. W. Hockney, “Parametrization of computer performance,” Parallel Computing, vol. 5, no. 1-2, pp. 97–103, 1987.

39. N. Dierckxsens, T. Li, J. R. Vermeesch, and Z. Xie, “A benchmark of structural variation detection by long reads through a realistic simulated model,” Genome biology, vol. 22, no. 1, pp. 1–16, 2021.

40. B. Grüning, R. Dale, A. Sjödin, B. A. Chapman, J. Rowe, C. H. Tomkins-Tinch, R. Valieris, J. Köster, and B. Team, “Bioconda: sustainable and comprehensive software distribution for the life sciences,” Nature methods, vol. 15, no. 7, pp. 475–476, 2018.

41. P. Biosciences, “PacBio Real-world HiFi long reads for O. sativa.” https://downloads.pacbcloud.com/public/dataset/Sequel-IIe-202104/rice/, 2021 (xlast date accessed: Aug 2022).

42. D. A. Benson, M. Cavanaugh, K. Clark, I. Karsch-Mizrachi, D. J. Lipman, J. Ostell, and E. W. Sayers, “Genbank,” Nucleic acids research, vol. 41, no. D1, pp. D36–D42, 2012.

43. W. Huang, L. Li, J. R. Myers, and G. T. Marth, “Art: a next-generation sequencing read simulator,” Bioinformatics, vol. 28, no. 4, pp. 593–594, 2012.

44. S. F. Altschul, W. Gish, W. Miller, E. W. Myers, and D. J. Lipman, “Basic local alignment search tool,” Journal of molecular biology, vol. 215, no. 3, pp. 403–410, 1990.

